# Immature human engineered heart tissues engraft in a guinea pig chronic injury model

**DOI:** 10.1101/2022.07.07.499077

**Authors:** Constantin von Bibra, Aya Shibamiya, Andrea Bähr, Birgit Geertz, Maria Köhne, Tim Stuedemann, Jutta Starbatty, Nadja Hornaschewitz, Xinghai Li, Eckhard Wolf, Nikolai Klymiuk, Markus Krane, Christian Kupatt, Bernhard Hiebl, Thomas Eschenhagen, Florian Weinberger

## Abstract

Engineered heart tissue (EHT) transplantation represents an innovative, regenerative approach for heart failure patients. Late preclinical trials are underway, and the first clinical trial has started in 2021. Preceding studies revealed functional recovery after implantation of in vitro-matured EHT in the subacute stage while transplantation in a chronic injury setting was less efficient. We hypothesized that the use of immature EHT patches (EHT^Im^) could improve cardiomyocytes (CM) engraftment. Chronic myocardial injury was induced in a guinea pig model (n=14). EHT^Im^ (15×10^6^ cells) were transplanted directly after casting. Functional consequences were assessed by serial echocardiography. Animals were sacrificed four weeks after transplantation and hearts were excised for histological analysis. Cryo-injury lead to large transmural scars amounting to 26% of the left ventricle. Grafts were identified by a positive staining for human Ku80 and dystrophin, remuscularizing 9% of the scar area on average. The CM density in the graft was higher compared to previous studies with in vitro-matured EHTs and showed a greater population of immature CM. Echocardiographic analysis showed a small improvement of left ventricular function after EHT^Im^ transplantation. In a small translational proof-of-concept study human scale EHT^Im^ patches (4.5×10^8^ cells) were epicardially implanted on healthy pig hearts (n=2). In summary, we provide evidence that transplantation of immature EHT patches without pre-cultivation results in better cell engraftment.

## Introduction

Heart failure is still an unsolved problem after myocardial injury^1^. Myocardial injury is followed by a remodeling cascade aiming to compensate for the irreversible loss of cardiomyocytes. Current therapeutic approaches are not able to address the loss of myocytes. New therapeutic approaches are therefore required^2^. Regenerative therapies have been successfully applied in preclinical models^3–7^. Transplantation of engineered cardiac constructs, e.g. EHTs has become a meaningful attempt. For this strategy, human induced pluripotent stem cell (hiPSC) derived CMs are embedded in a three-dimensional matrix and cultured over several weeks before transplantation. During the culture period CM mature considerably, coherently start to beat and at the time of transplantation resemble human myocardium, both morphologically (e.g. cell alignment and sarcomeric organization) and physiologically (e.g. action potential characteristics, response to pharmacological stimuli) ^4,8,9^. EHT transplantation remuscularized the injured heart in a dose-dependent manner and improved left-ventricular function in the highest dose^4^. The data indicate that a substantial part of the scar must be remuscularized for a meaningful effect. Yet, the beneficial effect was only seen in a subacute setting. EHT transplantation in the clinically more relevant chronic injury model was less efficient^10,11^, both in terms of graft formation and functional recovery^12^. While analyzing graft structure at early time points after transplantation, we noticed a temporary deterioration in CM structure. Over time, CM structure recovered and the cells formed human grafts with advanced cell organization. We therefore hypothesized that the transplantation of non-contracting immature EHT patches (EHT^Im^) can improve transplantation success in the more challenging chronic setting.

## Materials and Methods

### Cardiac differentiation and generation of immature EHT

Cardiomyocytes were differentiated from hiPSCs (UKEi1-A) as previously described^4^. UKEi001-A was reprogrammed by a Sendai Virus (CytoTune iPS Sendai Reprogramming Kit, ThermoFisher). EHT patch casting was performed by mixing CMs and a fibrinogen/thrombin mix^3^. EHT patches (1.5×2.5 cm) were generated with 15×10^6^ hiPSC CM per 1.5 to 1.6 ml for the guinea pig study. For the pig study large, human scale EHT patches (5.0×7.0 cm) with 4.1 and 4.8×10^8^ cells were cast. Cells were resuspended in 25 ml. All EHT^Im^ patches were subsequently transplanted within 2-5 hours.

### Animal Care and Experimental Protocol Approval

The investigation was performed in accordance with the guide for the care and use of laboratory animals published by the National Institutes of Health (Publication No. 85-23, revised 1985) and was approved by the local authorities (Behörde für Gesundheit und Verbraucherschutz, Freie und Hansestadt Hamburg: 109/16 and Regierung von Oberbayern: 02-18-134).

### Guinea pig chronic injury model and immature EHT transplantation

Female Dunkin Hartley guinea pigs were used (400-460 g, 6-8 weeks of age, Envigo) to induce chronic myocardial injury^12^. Cryo-injury of the left ventricular wall was performed with a liquid nitrogen cooled probe. Four weeks after injury a re-thoracotomy was performed and the cryo-lesion was covered with an EHT^Im^ patch (15×10^6^ hiPSC-CM) or a cell-free patch. The study was performed in parallel to a previous study in which matured EHTs were transplanted^12^. Animals that received cell-free patches served as control group for both studies. Guinea pigs were immunosuppressed beginning two days prior to transplantation with cyclosporine (7.5 mg/kg for the first five postoperative days and 5 mg/kg per day over the follow up period of four weeks; mean plasma concentration: 562±86.9 μg/l; n=7) and methylprednisolone (2 mg/kg).

### Echocardiography

Cardiac function was assessed by transthoracic echocardiography^13^ with a Vevo 3100 System (Fujifilm VisualSonics). Echocardiographic recordings were taken at baseline, four weeks after cryo-injury and four weeks after transplantation. Because of technical issues with the system three animals did not received baseline echocardiography. One animal was excluded from the functional analysis, because of inadequate image quality.

### Heart preparation and histology

Four weeks after transplantation hearts were processed for histology^4^. Hearts were fixed in formalin for two days, sliced into four cross-sections and paraffin-embedded for histology. Images of dystrophin and human Ku80 stained slides were acquired with a Hamamatsu Nanozoomer whole slide scanner and analyzed with NDP software (NDP.view 2.3.1). Dystrophin stained sections were used to determine infarct size and graft area^3,4^. Grafts were recognized by human Ku80 staining. To assess CM density, human Ku80^+^ nuclei were automatically counted (Q-Path 0.3.0) in 200 μm x 200 μm fields. Human Ku80^+^ nuclei quantification was performed in the section that contained the largest graft. The primary antibodies used are listed in the Supplemental Table. Immunofluorescence images were acquired with an LSM 800 confocal microscope (Zeiss).

### Immature EHT transplantation onto healthy pig heart

Transgenic pigs overexpressing a human CTLA4-Ig derivate (LEA29Y) were used for the transplantation study^14^. LEA29Y pigs were anesthetized by intramuscular injection of ketamine, azaperone and atropine sulfate and mechanically ventilated. Anesthesia was maintained by a constant intravenous application of propofol and fentanyl. EHT patches were transplanted through a left-sided lateral thoracotomy^4^. One large, human-scale EHT^Im^ patch per heart was sutured onto the healthy epicardium. LEA29Y pigs were additionally pharmacologically immunosuppressed with methylprednisolone (250 mg on the day of surgery and 125 mg/d starting at day 1 after surgery), tacrolimus (0.2 mg/kg per day), and mycophenolate mofetil (40 mg/kg per day). Animals were sacrificed under anesthesia by intravenous injection of saturated potassium chloride (3 days and 14 days after transplantation).

### Statistical Analysis

Statistical analyses were performed with GraphPad Prism 8.3.0 or 9.3.1, USA. Comparison among two groups was made by two-tailed unpaired Student’s t-test. One-way ANOVA followed by Tukey’s-Test for multiple comparisons were used for more than 2 groups. When two factors affected the result (e.g. time point and group), two-way ANOVA analyses and Tukey’s-Test for multiple comparisons were performed. Error bars indicate SEM. P-values are displayed graphically as follows: *p<0.05, ****p<0.0001.

## Results

### Generation of immature EHT patches

In this study we examined whether transplantation of EHT^Im^ patches could enhance CM engraftment in a chronic injury setting. In accordance with previous studies, EHT patches were cast with 15×10^6^ cells^4,12^ (troponin t positivity 94±1.8%; n=7). Whereas EHT patches were cultured for 21 day in previous studies, they were used for transplantation directly after casting in the current study. EHT patches remodel during the culture period, i.e. EHT^Im^ patches were slightly larger. Cell density in the patch was lower and the CMs less mature (Supplemental Figure 1). Human-scale EHT patches for the pig experiments were casted with 4.1 and 4.8×10^8^ cells, respectively (troponin t positivity 91.2%; n=2) and similarly transplanted within 12 hours after casting.

### Engraftment of immature EHT

Four weeks after the induction of myocardial damage, a re-thoracotomy was performed and the scar was covered with an EHT^Im^ patch or cell-free patch (n = 7, respectively). After a follow up period of four weeks, hearts were excised and prepared for histological analysis (Figure 1A and B). Histomorphometry exhibited similar scar areas in the cell-free control group (28.6±3.7% of the left ventricle, n = 7) and the intervention group (22.6±2.1%; n = 7; Figure 1C). Human origin of the grafts was demonstrated by human Ku80 immunostaining (Figure 1B). Six out of seven hearts in the EHT^Im^ group contained engrafted CMs. The graft area amounted to 9.4±3.9% of the total scar area (Figure 1D). The number of engrafted cells amounted to 2699±857 human cells per section (Figure 1E).

**Figure 1.**
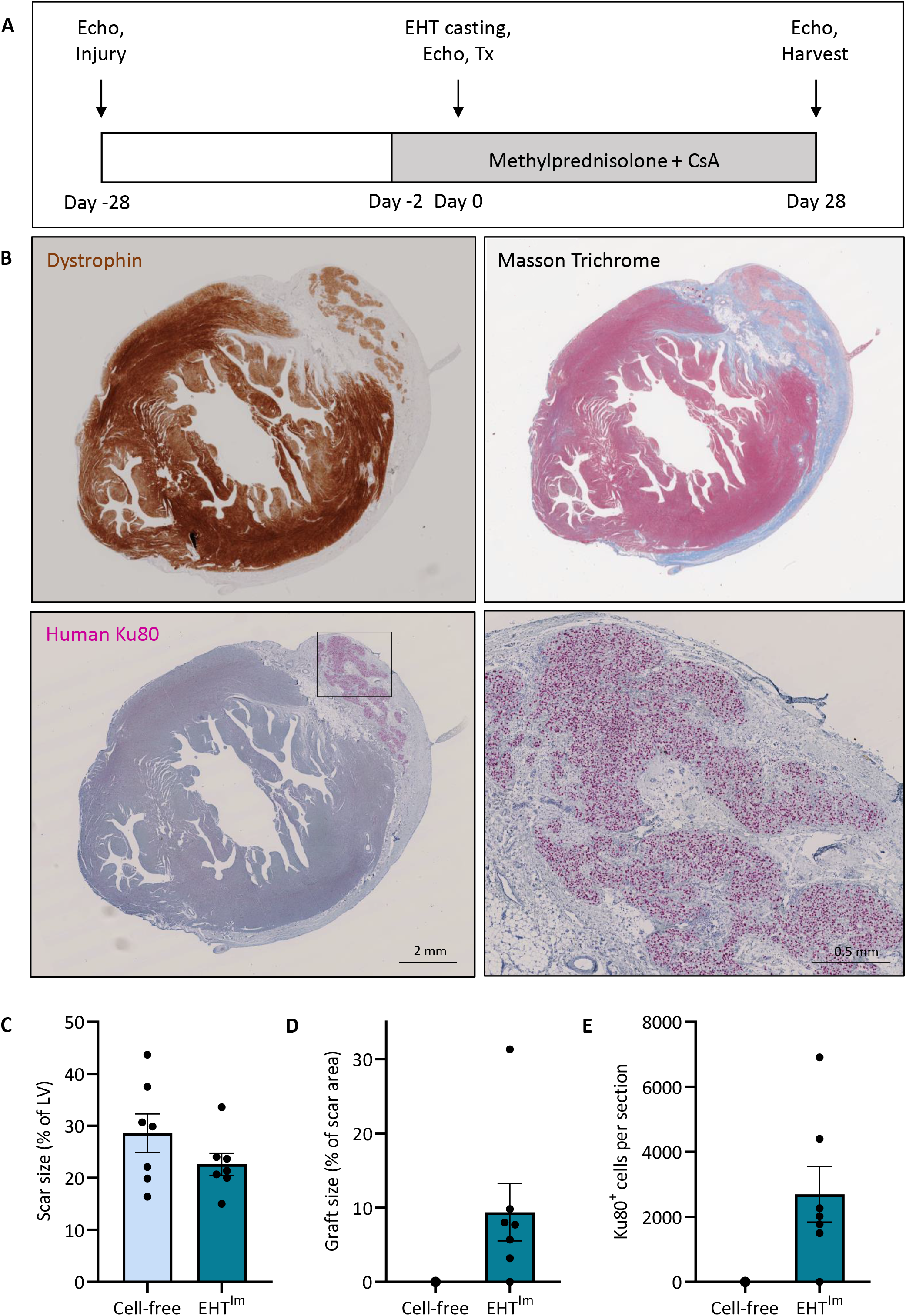
Histological assessment of guinea pig hearts after EHT^Im^ transplantation. **A**) Study design. **B**) Sections of one heart four weeks after transplantation stained for dystrophin, Masson Trichrome and human Ku80, graft shown in higher magnification. **C**) Quantification of scar size and (**D**) graft size. **E**) Quantification of Ku80^+^ cells per section. Each data point represents one heart. Mean ± SEM values are shown. Echo indicates Echocardiography; Tx, transplantation; CsA, Cyclosporine A; EHT^Im^, immature engineered heart tissue; LV, left ventricle.

Human grafts consisted of densely packed CM with advanced sarcomeric structure (sarcomere length 1.8±0.0 μm; n = 3; Figure 2A and D). In some sections, single engrafted human CMs with advanced morphologically structure were visible in the surrounding scar tissue, without apparent connection to the clustered graft (Supplemental Figure 1D). A subpopulation of the engrafted CM expressed the immature/fetal atrial isoform of myosin light chain (MLC2a 13.3±0.9%; Figure 2B and E), whereas most cells expressed the ventricular isoform (MLCC2v 86.7±0.9%), indicating maturation after transplantation. In contrast to this finding, most cells still expressed the slow skeletal troponin I isoform (ssTnI 89.4±1.2%; Figure 2C and F). Cardiac troponin I was weakly expressed (cTnI 10.6±1.2%). Additional evidence of immaturity was the circumferential N-cadherin expression (Figure 3A) and the ongoing cell cycle activity. Four weeks after transplantation, 9.7±1.3% of CM were still in the cell cycle as indicated by the expression of Ki67 (Figure 3D and E). Vascularity in the graft was lower than in the remote myocardium (368±17 vessels per mm^2^ vs. 1525±50 vessels per mm^2^ in the host myocardium; n=3, respectively; Figure 3B and C).

**Figure 2.**
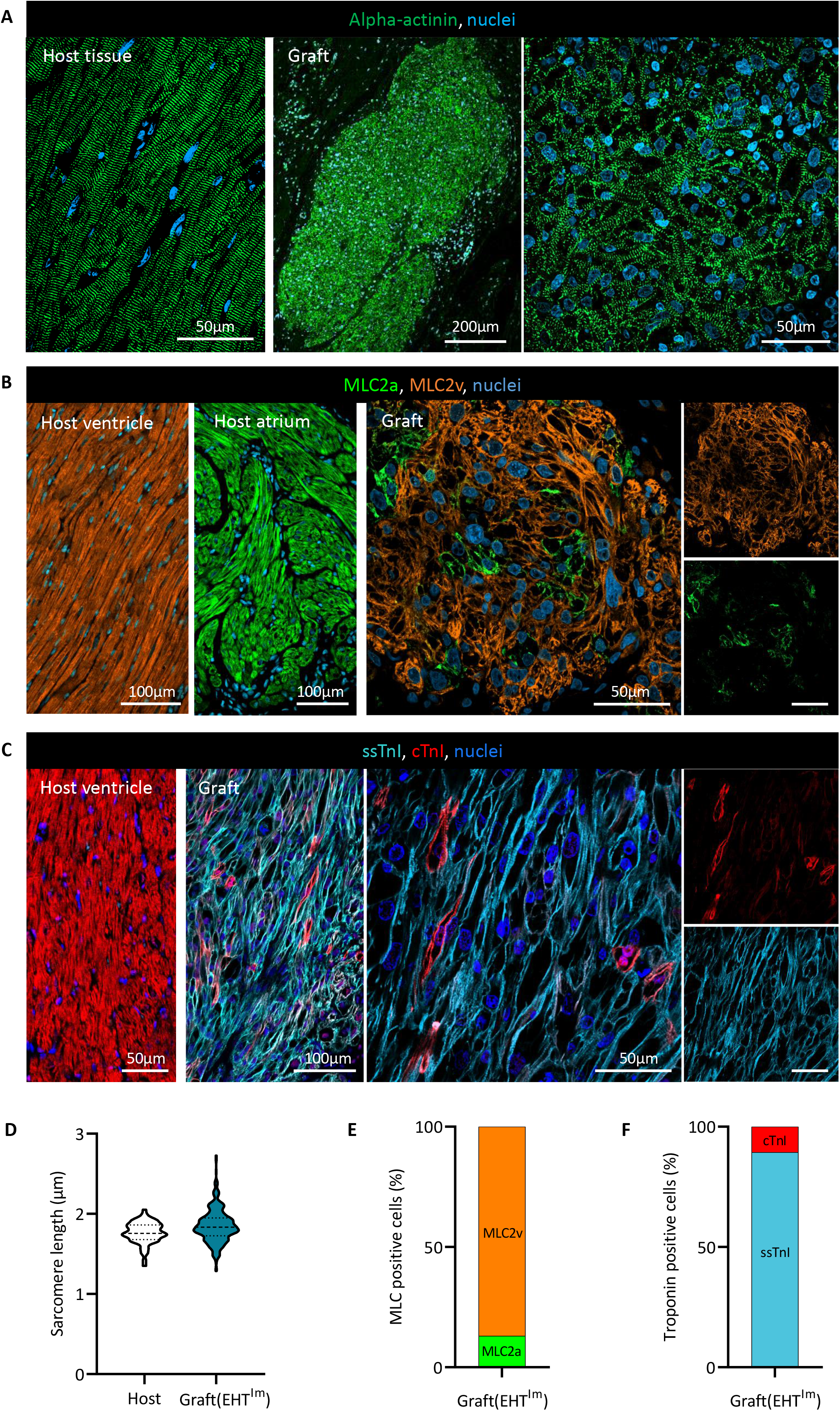
Structural analysis of engrafted cardiomyocytes. **A**) Alpha-actinin, **B**) myosin light chain (MLC2) and **C**) troponin I (TnI) isoform staining of host and graft myocardium. **D**) Quantification of sarcomere length in human grafts compared to guinea pig host myocardium in alpha-actinin stained sections (n=3 hearts/>90 sarcomeres per graft or host myocardium). **E**) Quantification of the atrial (MLC2a) and ventricular (MLC2v) isoform expression. **F**) Quantification of the slow skeletal (ssTnI) and cardiac troponin I (cTnI) isoform expression. EHT^Im^ indicates immature engineered heart tissue.

**Figure 3.**
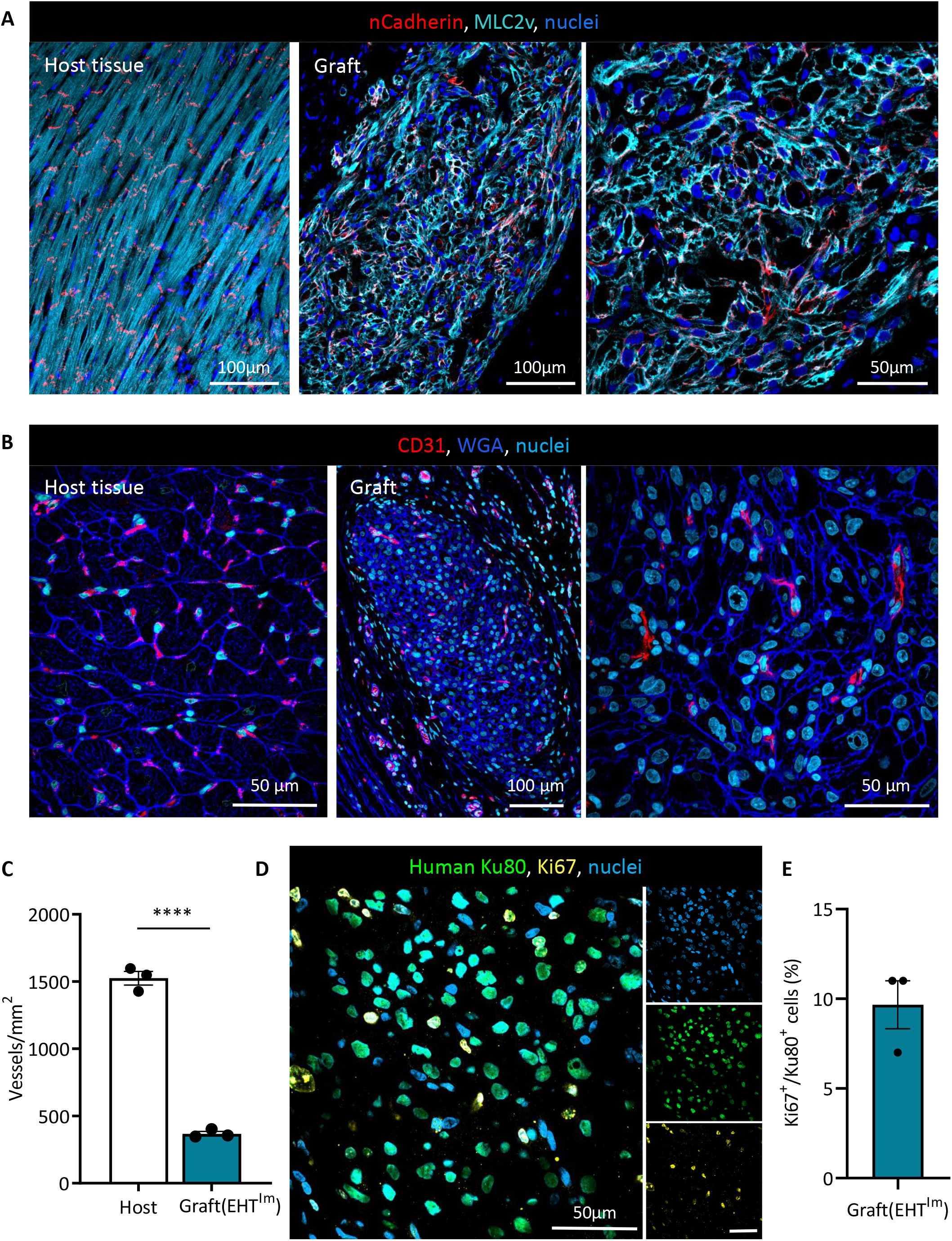
Analysis of vascularization and graft proliferation. **A**) N-cadherin and myosin light chain, ventricular isoform (MLC2v) staining of host and graft tissue. **B**) Human graft and host myocardium stained for WGA (wheat germ agglutinin) and CD31 and **C**) quantification of graft vascularization four weeks after transplantation (n=3 hearts/N=3 high magnification images per graft or host myocardium). **D**)Human Ku80/Ki67 double immunostaining of a human graft **E**) Ki67/Ku80 positive nuclei four weeks after transplantation (n=3 hearts/3 high magnification images per graft). Each data point represents one heart. Mean ± SEM values are shown, ****P<0.0001. EHT^Im^ indicates immature engineered heart tissue.

Additional graft analysis showed that cell density after transplantation of EHT^Im^ was significantly higher (cell density: 4809±507 Ku80^+^ cells per mm^2^; n=3) compared to grafts after the transplantation of in vitro-matured EHT patches^12^ (cell density: 3233±128 Ku80^+^cells per mm^2^; n=3; Figure 4A). To assess whether the higher density was related to a smaller cell size, cell surface area was evaluated by WGA (wheat germ agglutinin) and human Ku80 staining. Cell size was not smaller four weeks after transplantation (EHT^Im^: 109.1±2.2 μm^2^ vs. EHT: 90.4±1.8 μm^2^; n=3, respectively; Figure 4B).

**Figure 4.**
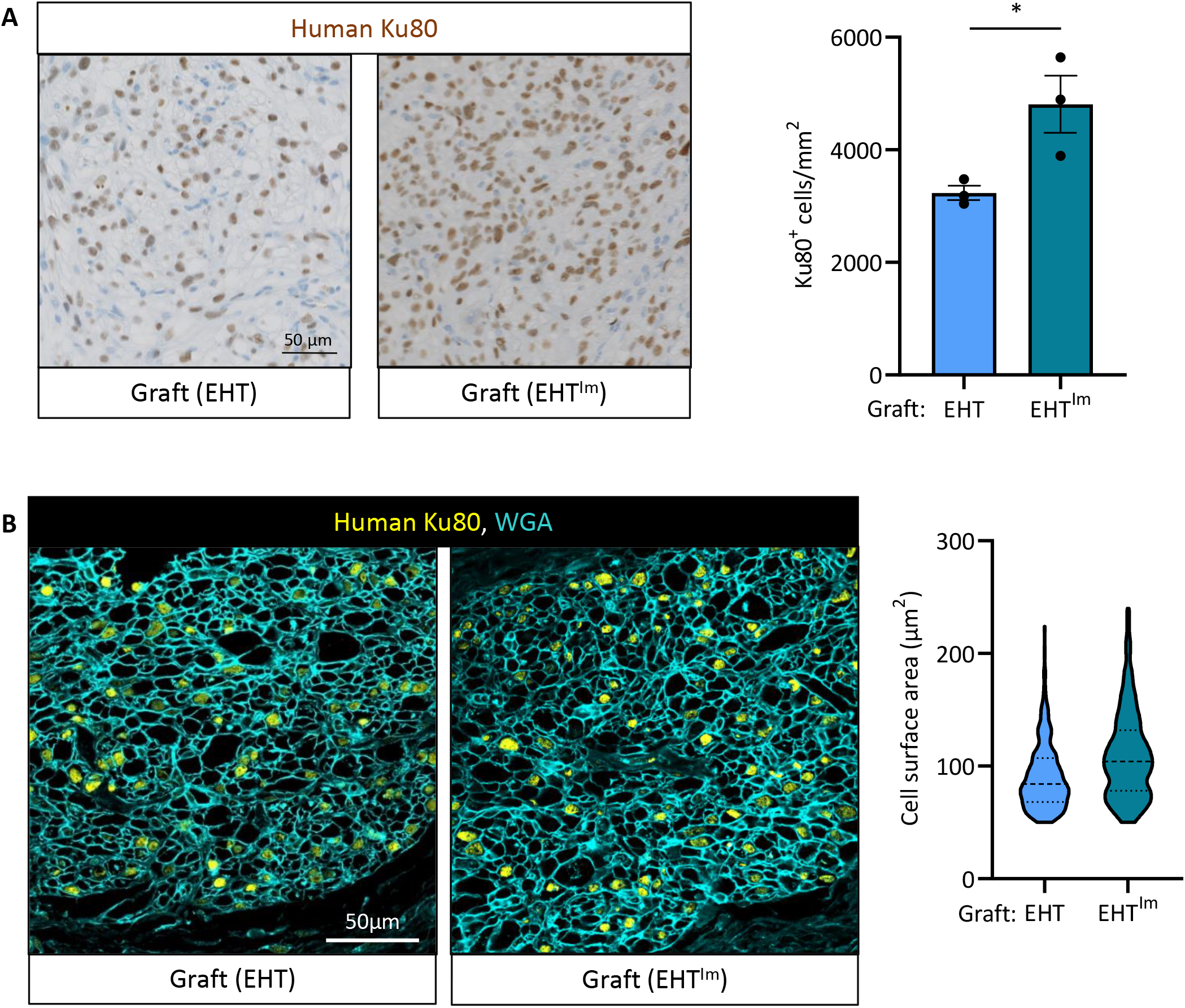
Comparison of engrafted EHT and EHT^Im^. **A**) Human Ku80 staining of a human graft four weeks after transplantation of an EHT or EHT^Im^ respectively and quantification of the Ku80^+^ nuclei per mm^2^ (n=3 hearts/3 high magnification images per graft). **B**) Human Ku80 and WGA (wheat germ agglutinin) staining of an EHT^Im^ four weeks after transplantation and quantification of cell size of human Ku80^+^ cells (n=3 hearts/N=3 high magnification images per graft, respectively). Each data point represents one heart. Mean ± SEM are shown, *P<0.05. EHT indicates engineered heart tissue; EHT^Im^, immature engineered heart tissue.

### Functional effects on left ventricular function

Functional effects of EHT^Im^ patch transplantation were assessed by serial echocardiography (Figure 5). Fractional area change (FAC) at baseline was 46.0±2.8% and fractional shortening (FS) was 50.6±2.0%. Myocardial injury resulted in a reduction of left ventricular function four weeks after injury (FAC: control group: 30.7±3.5% vs. EHT^Im^ group: 33.5±2.5%; FS: control group: 37.9±1.9% vs. EHT^Im^ group: 36.0±3.0%). FAC showed no improvement in left ventricular function (FAC: control group: 33.7±3.3% vs. EHT^Im^ group: 36.9±4.2%) whereas FS was higher in the EHT^Im^ group compared to animals that received cell-free patches (FS: control group: 34.1±3.0% vs EHT^Im^ group: 45.5±4.2%).

**Figure 5.**
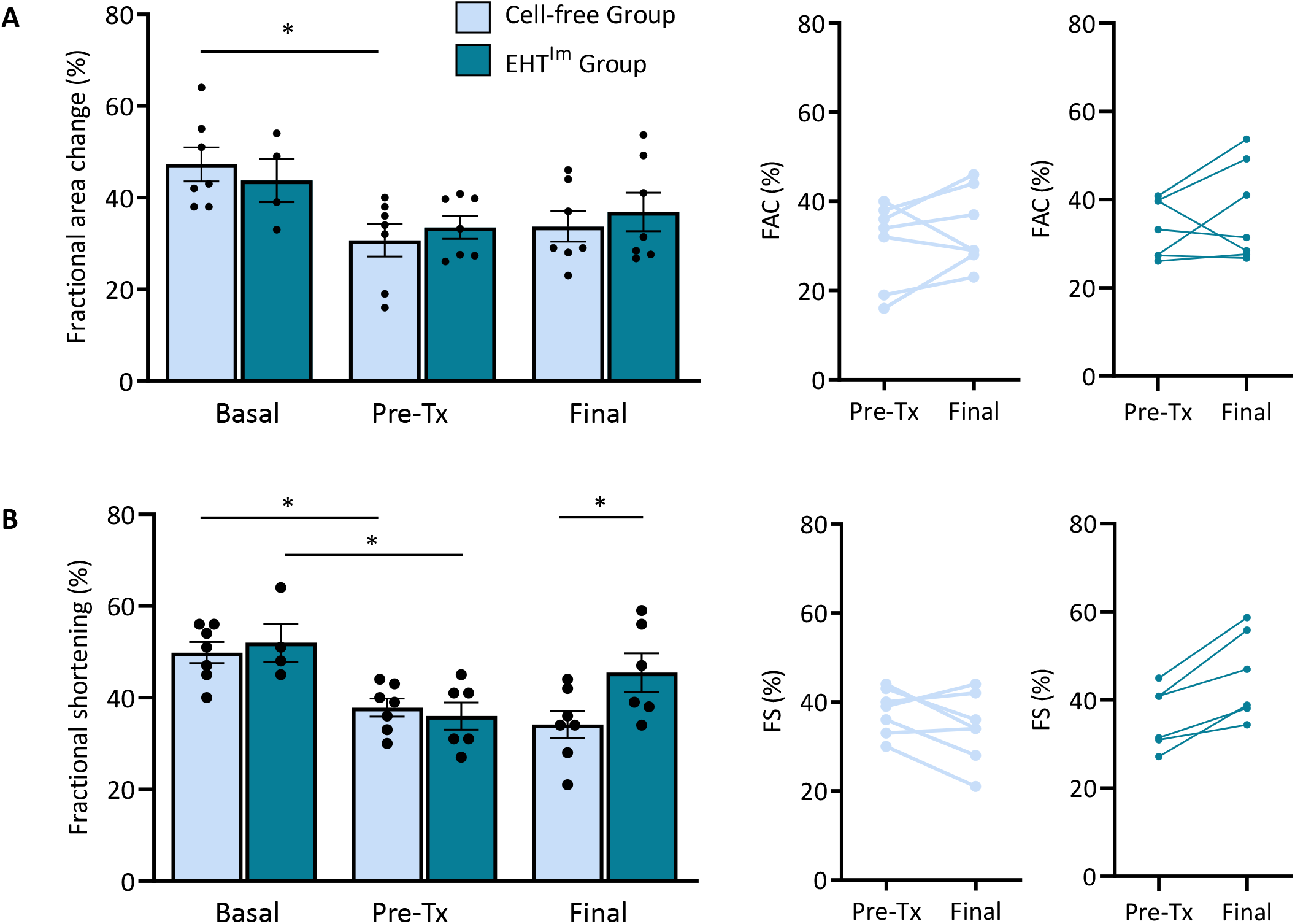
Echocardiographic evaluation. **A**) Fractional area change (FAC), **B**) fractional shortening (FS) and **C**) ejection fraction (EF) analysis at baseline, four weeks after injury and four weeks after transplantation (n=7 cell-free group, n=6 EHT^Im^ patch group) and differences between day 28 (post injury) and four weeks after transplantation of the respective groups. Mean ± SEM are shown, *P<0.05. EHT^Im^ indicates immature engineered heart tissue, Pre-Tx, before transplantation (post injury).

### Transplantation of large immature EHT in a pig

As a further translational step, large immature EHT patches were generated and transplanted on healthy pig hearts (5×7 cm, 4.1–4.8×10^8^ cells; n=2 EHT^Im^ patches). Human scale EHT^Im^ patches were epicardially transplanted in LEA29Y pigs (n = 2). Animals were sacrificed 3 days and 14 days after transplantation (Figure 6A). Histological analysis three days after transplantation showed large number of human CMs, mainly localized in the former surface parts of the EHT^Im^ patch (Figure 6B). Sarcomeric organization was low and no invasion of vessels was detected in the early days after transplantation (Figure 6C). Two weeks after transplantation, small grafts were visible (Figure 6D). Cardiomyocyte structure and vascularization was more advanced compared to the three days timepoint (Figure 6E).

**Figure 6.**
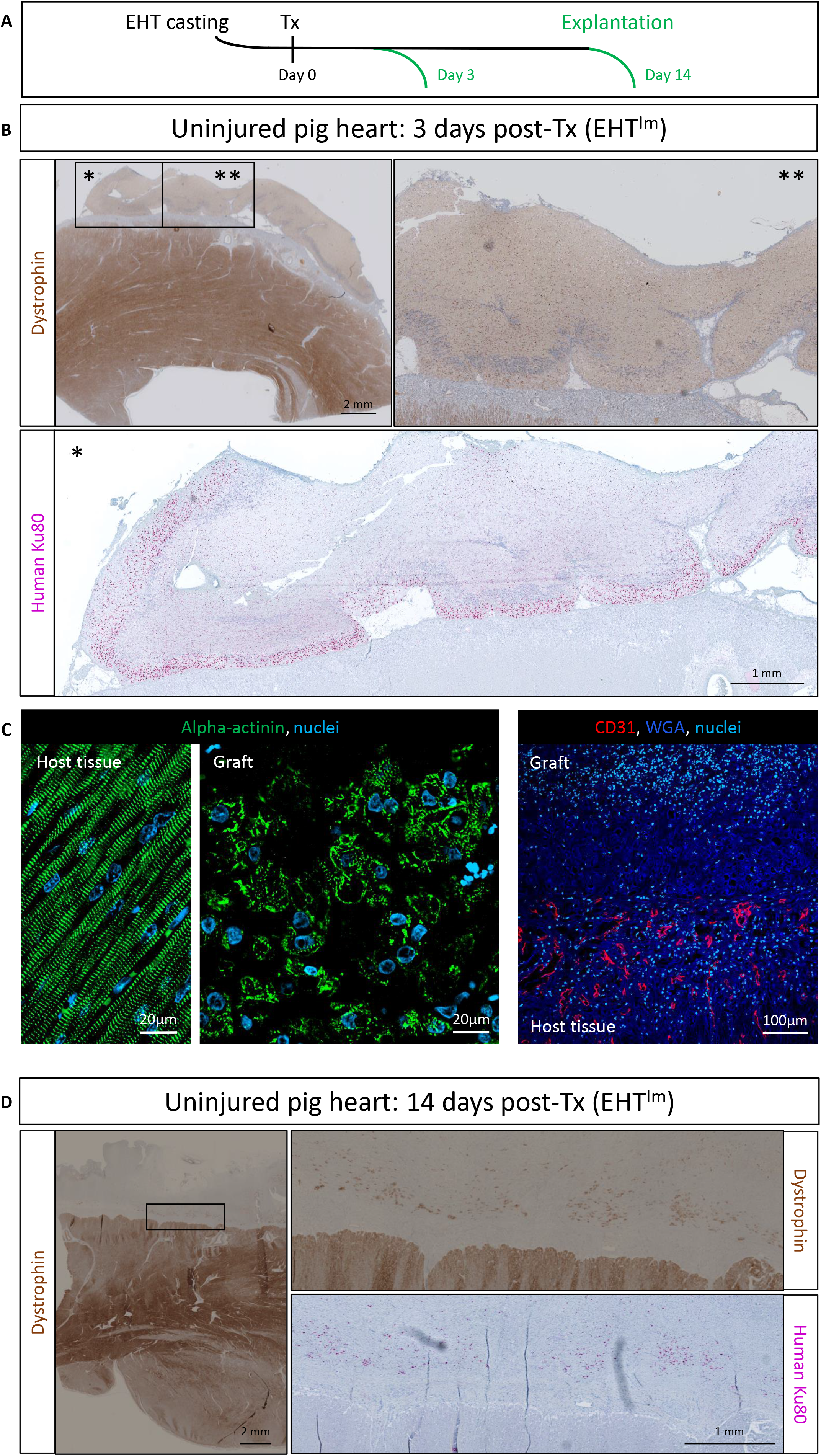
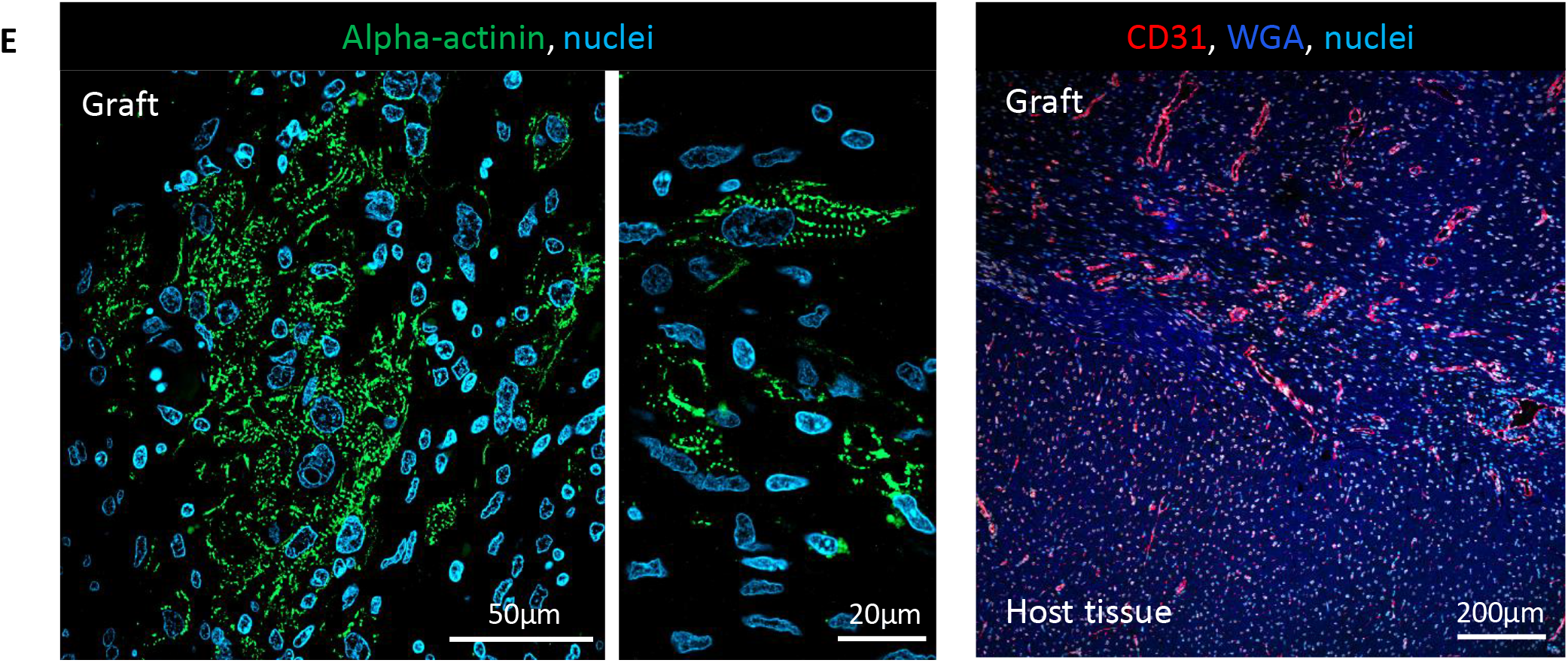
Histological evaluation after EHT^Im^ transplantation in pigs. **A**) Experimental protocol. **B**) Sections of one heart three days after transplantation stained for dystrophin and human Ku80, graft shown in higher magnification. **C**) Alpha-actinin staining for host and graft myocardium and staining for WGA (wheat germ agglutinin) and CD31. **D**) Sections of one heart two weeks after transplantation stained for dystrophin and human Ku80, graft shown in higher magnification. **E**) Alpha-actinin staining for host and graft myocardium and staining for WGA (wheat germ agglutinin) and CD31. Tx indicates transplantation; EHT^Im^, immature engineered heart tissue.

## Discussion

In this study we investigated the transplantation of immature EHT patches in a guinea pig chronic injury model. Transplantation of hiPSC-derived CMs represents a promising conceptually new therapeutic strategy for heart failure patients^2^. First clinical trials have commenced in 2021^15,16^. The clinical trials are based on preclinical studies that repeatedly demonstrated that CM transplantation can improve, or at least stabilize, heart function after injury^3–5,17–24^. Yet, most of these studies were performed in subacute injury models, whereas most patients with advanced heart failure suffer from a chronic disease progression^25^. The few studies that evaluated CM transplantation in chronic injury models showed that CM engraftment in the chronically injured heart was less efficient. In a recent study EHT patch transplantation resulted in a lower degree of remuscularization when targeting the chronically injured heart^12^ compared to a transplantation in a subacute situation^4^. Likewise, other studies showed that the regeneration of a chronic injured heart^10,11,26^ or spinal cord^27,28^ is more challenging, indicating that this is not peculiarity of EHT transplantation but a more general problem for regenerative medicine.

Here we evaluated whether immature EHTs, i.e. EHT directly after casting, engraft better than matured EHTs with a coherent beating pattern. The study was based on the finding that transplanted matured EHTs showed remarkable dedifferentiation in the first week after transplantation on the injured heart^4^. At the time of transplantation, CMs in the matured EHTs morphologically resembled mature CMs (e.g. elongated cell shape, regular sarcomeric structure), but during the first days after transplantation most of this structure was lost and only after one to two weeks the surviving CMs recovered, restructured and regained a regular sarcomeric structure again. We therefore hypothesized that transplantation of more immature CMs in a three-dimensional tissue will circumvent this step and thereby improve engraftment.

The main differences after the transplantation of immature compared to matured EHT transplantation were: i) graft size was larger (albeit only to a small degree), ii) the engrafted CM were more immature, iii) cell density in the graft was higher and iv) FS showed an improvement in left-ventricular function. There was no difference in scar size, sarcomeric structure, cell cycle activity and graft vascularization after four weeks. Factors that might have contributed to better CM engraftment are: i) increased cell survival either via a faster vascularization or a higher resistance against hypoxia and mechanical stress, ii) a higher proliferation rate of the immature CM early after transplantation. Even though the total graft size was only slightly larger, the number of engrafted cells was significantly higher. This finding was surprising, in particular as the CMs were not substantially smaller. While the findings of this study are in line with work on rodent CMs demonstrating that neonatal (immature) CMs survive transplantation whereas adult ones do not^29^, they somewhat contradict own results that showed a more favorable effect when transplanting more mature EHTs (cultured in the presence of serum) over less mature EHTs that were cultured without serum^4^. Retrospectively, the stronger force of EHTs that were cultured in the presence of serum might not only have reflected maturity but overall cell wellbeing. The results of the present study also stand in contrast to recent CM injection studies in which CMs that were matured prior to transplantation engrafted better^30^, indicating either differences regarding the optimal cell maturity between the EHT transplantation and cell injection or that the optimal maturation state of the patch approach lies somewhere in between the recent and the previous study.

A small proof-of-concept study was performed in pigs to evaluate the feasibility of transplanting large EHT^Im^ in a pig model. A previous study revealed technical problems when transplanting smaller matured EHT patches^4^. Similar to matured human scale EHT patches, large EHT^Im^ patches resisted the forces of a pig heart and could be transplanted successfully. Three days after transplantation the patch was adapted to the epicardial surface but surprisingly not vascularized. Two weeks after transplantation, small vascularized grafts were formed.

Limitations: Cardiomyocyte purity of the EHT^Im^ was higher than in the previous study (troponin t positivity 94±1.8% vs. 76 ± 2.6%). This might have influenced the results, but in our experience cannot explain the difference between the two studies. Additionally, we cannot rule out that the number of transplanted CMs was different because cells were counted at the time of EHT generation and might have changed (diminished) over the culture period. Data from smaller EHTs indicate that cell count remains stable over time, but a slight difference might have gone unnoticed^31^. Echocardiographic analysis demonstrated an improvement in FS but not in FAC. This probably highlights the technical difficulties in performing echocardiography in this animal model after repeated thoracotomies. Additionally, anterolateral injuries as induced in our model can be underestimated by short-axis M-mode imaging^32,33^.

In summary, this study showed that transplantation of freshly prepared, immature EHT patches is feasible and remuscularizes the chronically injured heart at least as well as mature EHTs. The use of immature EHT comes at the cost of losing contractility of the patches as a central quality criterion, but would reduce practical challenges and costs of cultivating EHTs over extended time prior to transplantation.

## Acknowledgements

We thank Kristin Hartmann (UKE, Mouse Pathology Core Facility) for technical support with immunohistochemistry. We would also like to acknowledge the laboratory animal facility staff, UKE for their effort during the study.

## Author contributions

C.v.B. performed experiments, analyzed data and prepared the manuscript. A.S. performed cell differentiation and cast EHT patches. B.G. performed experiments (echocardiography and surgeries) and analyzed data. J.S., M.Ko., T.S. performed experiments (cell culture and histology). A.B., N.H., E.W., X.L., M.Kr., C.K. performed transplantation studies in pigs. N.K. provided LEA29Y pigs. B.H. supervised the project. T.E., F.W. designed and supervised the project, acquired funding and prepared the manuscript.

---

**Supplemental Figure 1.**
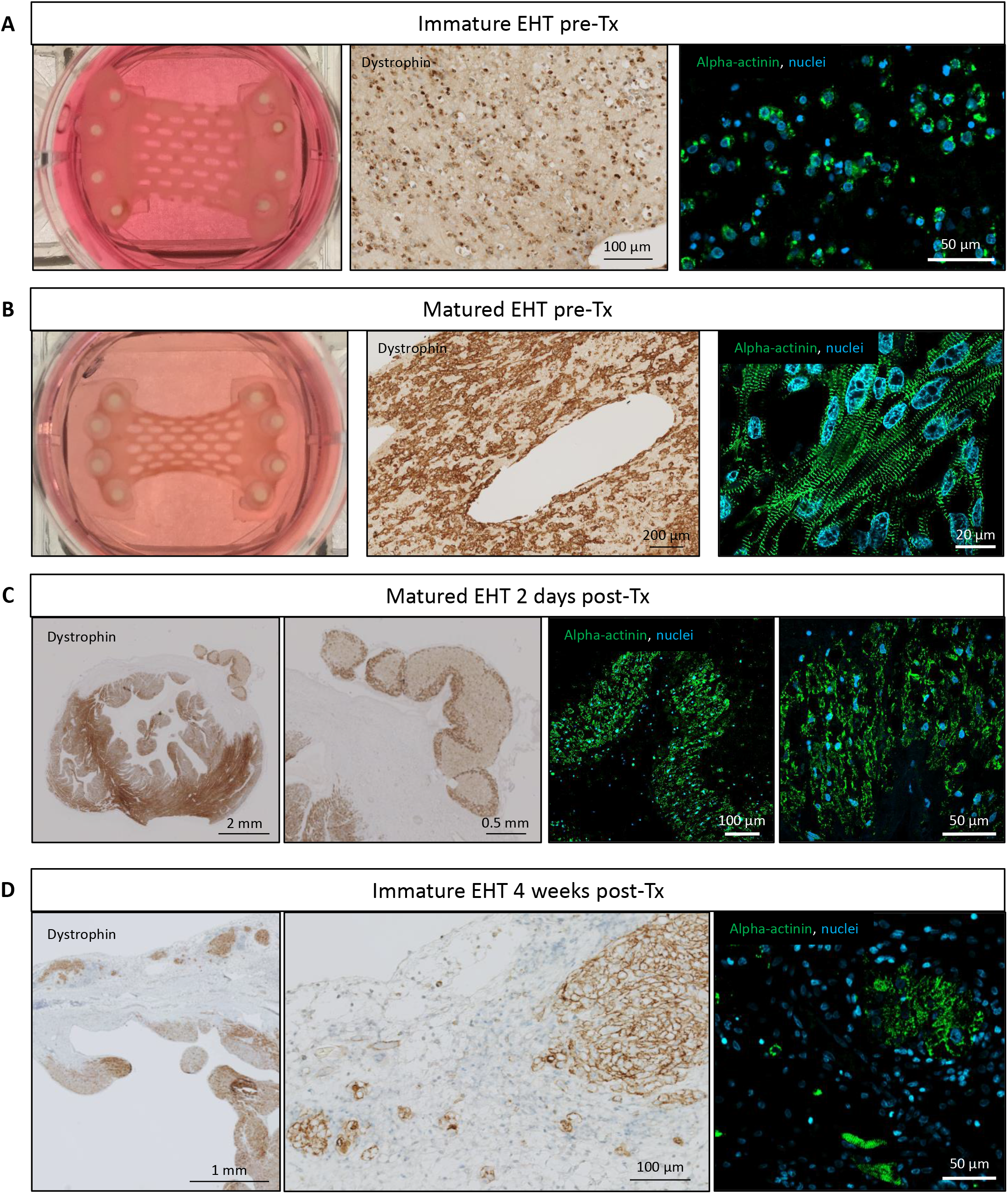
Development of engineered heart tissue. **A**) Photograph of an EHT at the time of casting (before transplantation) and dystrophin and alpha-actinin stained sections. **B**) Photograph of an EHT after 3 weeks in culture (before transplantation) and dystrophin and alpha-actinin stained sections. **C**) Dystrophin and alpha-actinin stained sections of an EHT heart 2 days after transplantation. **D**) Dystrophin and alpha-actinin stained sections of an EHT^Im^ heart 4 weeks after transplantation. EHT indicates engineered heart tissue. Tx, transplantation.

**Supplemental Table.** primary antibodies for histology.

| **Antigen** | **Vendor** | **Antigen retrieval** | **Titer** |
| --- | --- | --- | --- |
| **MLC2a** | BD Pharmingen, clone S58-205, Cat. 565496 | Citrate buffer | 1:500 |
| **MLC2v** | Proteintech, pAb, Cat.10906-1-AP | Citrate buffer | 1:250 |
| **α-actinin** | Sigma-Aldrich, clone EA-53, Cat. A7811 | Proteinase K | 1:600 |
| **Ku80** | Cell Signaling Technology, clone C48E7, Cat. CST-2180 | Citrate buffer | 1:800 |
| **N-cadherin** | Sigma-Aldrich, clone CH19, Cat. C1821 | Citrate buffer | 1:400 |
| **ssTnl** | Novus Biologicals, clone OT18H8 | Citrate buffer | 1:100 |
| **cTnl** | Abcam, Cat. Ab47003 | Citrate buffer | 1:100 |
| **PECAM1**<br>**(CD31)** | Novusbio, Cat. NB 100-2284 | Citrate buffer | 1:100 |
| **Ki67** | Invitrogen, clone Sol-A15, Cat. 2196793 | Citrate buffer | 1:400 |

